# evo3D R package: a spatial haplotype framework for structure-informed analysis of molecular evolution

**DOI:** 10.64898/2025.12.12.694041

**Authors:** Bradley K. Broyles, Qixin He

**Affiliations:** Department of Biological Sciences, Purdue University, West Lafayette, IN, USA; Purdue Institute of Inflammation, Immunology and Infectious Disease, West Lafayette, IN, USA

## Abstract

At the molecular level, selection pressures often act on protein structural features, yet most evolutionary analyses remain confined to linear sequences. Early structure-informed approaches improved interpretability by mapping single-site metrics onto protein structures, and later methods introduced 3D sliding windows to capture spatially clustered signals missed by linear window approaches. These frameworks, however, are restricted to predefined statistics and narrowly defined 3D window types, limiting the scope of questions that can be addressed. We developed an R package, `evo3D`, as a new framework for structure-informed evolutionary analysis that supports a wide range of downstream statistics and scales from simple to complex structures. `evo3D` extracts structure-informed multiple sequence alignment subsets (spatial haplotypes), making the structure-informed unit of analysis directly available to users. The framework supports fixed-count and fixed-distance spatial windows, introduces residue and codon analysis modes, and extends to multimers, interfaces, and multiple structural models through a single wrapper, `run_evo3d()`. We demonstrate `evo3D`’s utility by performing an epitope-level diversity scan of Hepatitis C virus E1/E2 complex, identifying conserved spatial neighbourhoods missed by linear sliding windows, and by evaluating `evo3D`’s scalability on the octameric Chikungunya virus E1/E2 assembly. Importantly, `evo3D` formalises the core components of structure-informed analysis of molecular evolution and removes technical barriers. As a result, the framework streamlines the evaluation of evolutionary patterns directly within 3D structural contexts, and we anticipate its wide application in molecular evolution studies. The package is available at github.com/bbroyle/evo3D.

## Introduction

Selection pressures acting on protein-coding regions frequently target three-dimensional (3D) features such as catalytic domains, ligand-binding sites, protein-protein interfaces, and overall structural stability, where the selected phenotypes emerge from neighbourhoods of spatially clustered residues (Golding and Dean 1998; Echave et al. 2016). Linear sequence approaches, whether based on single sites or sliding windows, cannot capture this spatial context and are therefore limited in inferring selection on structural features. Incorporating 3D information has improved detection of conserved interaction surfaces (Lichtarge et al. 1996; Armon et al. 2001; Landgraf et al. 2001; Chivot et al. 2025), interpretability of per-codon rates of evolution (Knight et al. 2007), and detection of adaptive evolution at epitopes and ligand-binding regions (Suzuki 2004; Berglund et al. 2005; Guy et al. 2018; Ciubotariu et al. 2024). Yet existing approaches for incorporating structural context remain restricted to fixed statistics and narrowly defined spatial window types, limiting the scope of evolutionary questions that can be asked. We developed `evo3D`, a spatial haplotype framework for structure-informed analysis that supports a wide range of downstream genetic statistics and scales from monomers to multimeric assemblies.

The general goal of structure-informed analysis is to evaluate genetic diversity within spatially defined windows, requiring two inputs: a nucleotide or amino acid multiple sequence alignment (MSA) and a corresponding protein structure (PDB). Spatial windows are conceptually similar to linear sliding windows, with residues grouped by three-dimensional proximity rather than sequential proximity. We define a spatial haplotype as the codon sequence formed by the subset of MSA columns corresponding to residues in a spatial window. Despite this conceptual similarity to linear sequence windows, constructing spatial windows in practice is considerably more complex. The residues included in a window depend on the distance metric used to define inter-residue proximity and on the chosen structural model, as conformational differences between models alter spatial neighbourhoods.

Extending structure-informed analysis to multimeric assemblies further complicates spatial windows and raises biological and analytical considerations that have not been systematically formalised. A single codon may map to multiple residues across equivalent chains, possibly producing duplicated codons within a spatial window. In these cases, we introduce two window modes to specify how duplicated codons are treated: “residue” mode retains duplicated codons, whereas “codon” mode reduces windows to unique codons. Whether to retain or deduplicate these copies depends on the downstream statistic, with spatial averaging of single-site metrics generally requiring deduplication. Additionally, `evo3D` supports two analysis modes that determine how spatial windows are assigned. “Residue” analysis mode creates a separate spatial window for every structural residue, enabling comparisons across conformational states such as open and closed enzyme forms. “Codon” analysis mode collapses residue-level windows for a codon into one codon-level window, merging distance and solvent-exposure information across chains and multiple structural models. Spatial windows in multimers may also span residues encoded by different genes, raising the question of whether spatial haplotypes should allow combining segments of different genes from different genomes or be restricted to true haplotypes/genotypes from the same genome. This choice depends on the statistic, with haplotype-based measures generally requiring true haplotypes. `evo3D` formalises these multimer considerations within a unified framework, allowing users to select biologically and statistically appropriate definitions for their analysis.

Even at the monomer level, several methodological and implementation decisions continue to limit structure-informed analysis. Most tools are designed for a single pre-defined statistic, such as local missense mutation density in `HotMAPS` (Tokheim et al. 2016) and spatially correlated single-site substitution rates in `CONSTRUCT` (Chivot et al. 2025) and therefore do not provide a general-purpose framework. Others, such as `BioStructMap` (Guy et al. 2018), prepackage multiple statistics including Tajima’s D (Tajima 1989) and nucleotide diversity (Nei 1987) but are not easily extended to new statistics. A more general framework, which has not been previously implemented, is to return spatial haplotypes to users for easy applications of downstream analysis.

Existing approaches also present practical barriers for broader adoption, since the methods are either external-dependency heavy (`BioStructMap`) or restricted to specific operating systems (`HotMAPS`, `CONSTRUCT`). Although these methods make valuable contributions and include specialised features we have not implemented in `evo3D`, such as `BioStructMap`’s handling of intron-exon splicing, `CONSTRUCT`’s optimisation of window-distance cutoffs and distance-weighted spatial averaging, and `HotMAPS`’ design for large cancer mutation datasets, there remains a clear need for easy-to-implement, general-purpose methods that link MSAs and structures.

Methodologically, these previous approaches also inherit two persistent limitations that are shared across structure-informed workflows. First, the alignment between MSA codons and PDB residues is often unexamined. Reference nucleotide sequences must be translated and aligned to structural chains, yet existing methods treat this mapping as implicitly correct and do not expose it to users. This allows alignment errors, particularly in proteins with unresolved regions, to propagate into downstream analyses. Although potentially affecting only a few windows with a mis-mapped codon, identifying which windows were affected and correcting them is critical for accurate analysis. `evo3D` addresses this by returning internal alignments and allowing restarts of the workflow after user-adjusted mappings. Second, spatial windows in existing methods are defined exclusively by distance cutoffs, producing variable-count windows that complicate comparisons across sites and proteins. `evo3D` constructs either fixed-distance or fixed-count windows, depending on the desired analysis. Further extending existing approaches, `evo3D` can extract functional regions, such as protein interfaces or other user-defined windows, as spatial haplotypes for analysis alongside structure-derived windows.

Together, `evo3D` formalises a transparent and accurate mapping from MSAs to protein structures, spanning monomer handling to higher-order protein complexes. The framework introduces several methodological advances: (i) distinguishing window modes for handling duplicated codons; (ii) assigning windows at residue or codon levels via analysis mode; (iii) formalising cross-gene window considerations (iv) returning spatial haplotypes, the structure-informed unit of analysis, to users for downstream statistics; (v) implementing fixed-count windows for consistent window sizes; (vi) exposing internal MSA to PDB mapping for inspection and correction; and (vii) extracting protein-protein interfaces as independent spatial haplotypes. These advances extend the evolutionary questions that can be addressed within a structure-informed framework.

An additional goal of evo3D is to reduce practical barriers to performing such analyses. The `evo3D` R package, with minimal dependencies and cross-platform, has a single function entry point, `run_evo3d()`, that generalises across diverse structural contexts. Below, we outline the package workflow and demonstrate its application through analyses of two viral protein complexes. Interestingly, `evo3D` identifies conserved antibody-accessible regions that are missed in linear sequence approaches of the Hepatitis C virus E1/E2 complex; it also reveals elevated diversity adjacent to, but not within, receptor-binding domains in the octameric Chikungunya virus E1/E2 assembly. It is worth noting that our method is not limited to viral proteins, and the underlying MSAs are not restricted to within-species comparisons. The framework can be applied to many comparative evolution-driven questions where incorporating structural context is beneficial, with sequence depth and protein structures defined by the user. With accurate deep-learning structural prediction now routine (Jumper et al. 2021; Lin et al. 2023; Abramson et al. 2024), structural information is no longer a bottleneck, enabling `evo3D` to support a broad range of evolutionary analyses to be performed directly within the 3D environments that shape protein evolution.

### Implementation

`evo3D` is implemented in R (R Core Team 2024) and provides a single entry point, `run_evo3d()`, which performs a complete structure-informed evolutionary analysis (Fig. 1). The package maintains minimal external dependencies, using `bio3d` (Grant et al. 2006) for MSA and PDB input file handling, `msa` (Bodenhofer et al. 2015) for alignment utilities, `seqinr` (Charif and Lobry 2007) for nucleotide to amino-acid translation, `pegas` (Paradis 2010) for population-genetic statistics, `jsonlite` (Ooms 2014) for storing outputs to file, and `Rcpp` (Eddelbuettel and François 2011; Eddelbuettel 2013; Eddelbuettel and Balamuta 2018; Eddelbuettel et al. 2024) for high-performance solvent-accessibility calculations. The full parameter space is described in the package documentation and Supplementary Methods (Table S2). Default parameters can be inspected via `show_evo3d_defaults()`and adjusted as arguments to `run_evo3d()`. General guidance on parameter combinations suited to different types of analyses is provided in Supplementary Methods (Table S3).

**Figure 1.**
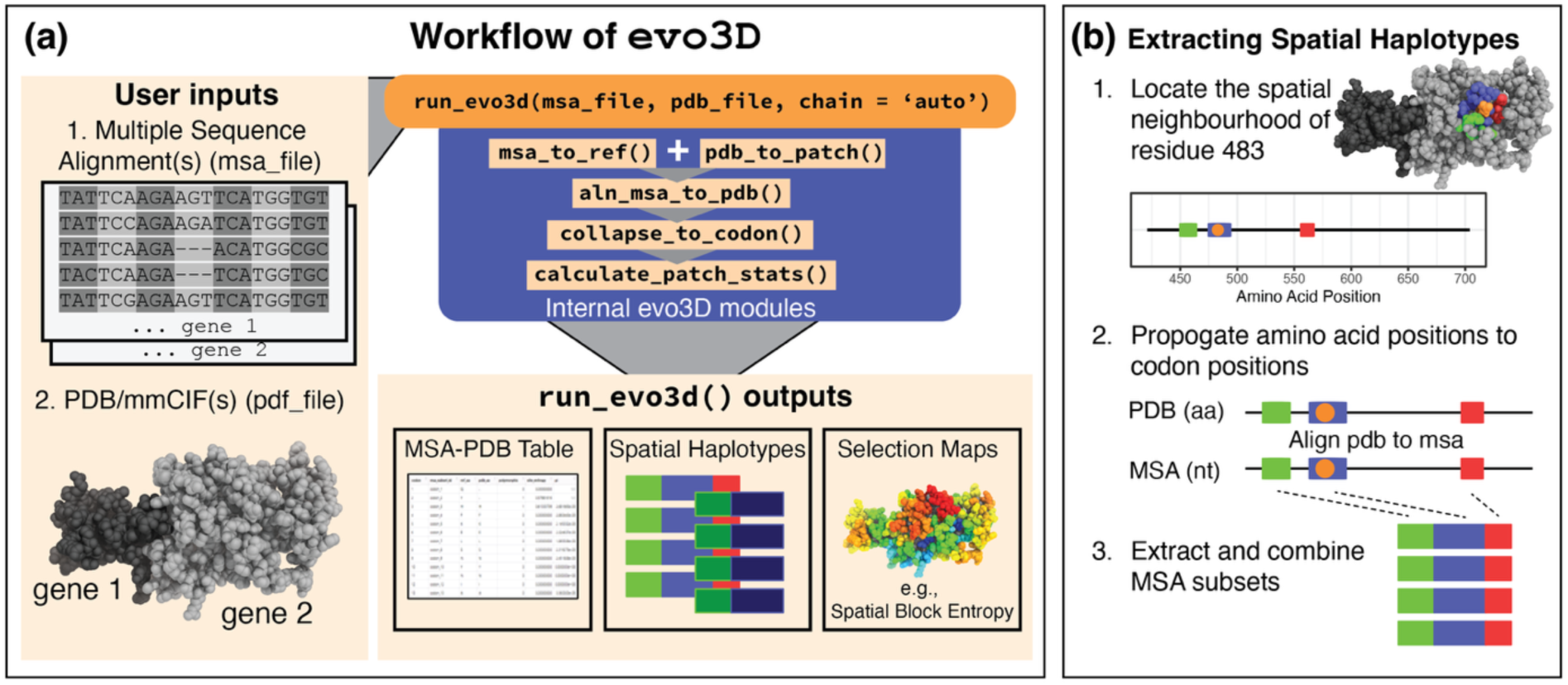
Illustration of the evo3D workflow and the spatial haplotype extraction process. **(a)** Required user inputs include a multiple sequence alignment (MSA) (nucleotide or amino acid) and a PDB/mmCIF structure (single chain or complex). In the example of user inputs, two gene MSAs correspond to one protein complex. `run_evo3d()` automatically maps structure chains to MSAs, defines 3D neighbourhoods (patches), aligns the PDB sequence to the MSA, and computes spatial haplotype statistics. Outputs include an MSA-PDB mapping table, spatial haplotypes (MSA subsets) (optionally written to file), and selection maps projected onto the structure. **(b)** Spatial haplotypes are constructed by (1) defining a 3D neighbourhood around a centroid residue (tuneable parameters control neighbourhood definition); (2) propagating residue positions to codon positions via the PDB→MSA map; and (3) extracting and concatenating those MSA columns to generate spatial haplotypes. The process is more complicated for multimeric assemblies (see the main text). *Abbreviations:* aa, amino acid; nt, nucleotide.

All results are returned as a structured S3 list containing:

- `evo3d_df` – the core results object, a data frame object with sequence-structure alignments, codon-indexed windows and window information, and computed statistics
- `final_msa_subsets` – list object of spatial haplotypes with names corresponding to the `msa_subset_id` column in `evo3d_df`
- `msa_info_sets` – input MSAs, internally generated reference sequence, translated reference sequence, and detected sequence type (nucleotide, protein)
- `pdb_info_sets` – input PDBs, `residue_df` with residue information and residue-indexed windows, peptide sequences for analysis chains, and residue-residue distance matrix
- `aln_info_sets` – `aln_df` with both codon and residue indexed windows, `coverage` for alignment quality, and `pos_mat`, which contains correctable MSA-PDB alignments.
- `call_info` – full parameter record, input file paths, and internal automatic chain assignments for reproducibility

The wrapper workflow begins with `msa_to_ref()`, which validates the MSA input, detects nucleotide versus amino-acid input, and generates a corresponding peptide reference sequence. Structural files are then loaded, and chains are matched to the MSA using fast k-mer similarity. Users may override this with `chain = “A“` to force specific mappings or chain strings such as `“ABC“` to force mappings to multiple chains. `pdb_to_patch()` defines spatial neighbourhoods around each residue using distance-based or fixed-count rules with user-set exposure and geometry parameters, producing residue-indexed 3D windows.

Next, `aln_msa_to_pdb()` aligns the reference peptide to the structural chain, maps MSA codons to residues, corrects gap-induced inconsistencies, and converts residue-indexed windows into codon-indexed windows. `collapse_to_codon()` merges per residue windows into per codon representations. `extract_msa_subsets()` isolates the MSA subsets corresponding to each spatial window, generating discontinuous but structurally coherent spatial haplotypes. Finally, `calculate_patch_stats()` applies the user-selected population-genetic or evolutionary statistics (e.g., block entropy (Olsen et al. 2011), nucleotide diversity (Nei 1987), haplotype diversity (Nei and Tajima 1981), Tajima’s D (Tajima 1989) to each haplotype.

The wrapper design minimises user workload, and tunable parameters are organized as control lists arguments (e.g., `pdb_controls = list(dist_cutoff = 10)`). All internal modules are exposed as standalone functions for custom workflows. Select submodular functions are exported with the package while others are accessible as internal functions (e.g. `evo3D:::.align_sequences()`). One of the useful submodular functions, `.calculate_accessibility()`, invoked by `pdb_to_patch()`,couples R and C++ to reproduce DSSP-style solvent-accessibility calculations (Kabsch and Sander 1983; Hekkelman et al. 2025). Additional utilities outside the wrapper function include functions that map statistics to PDB structures for visualisation, generate null distributions of synthetic linear/spatial windows that mirror the observed analysis codon inclusion and window definitions, and prune overlapping windows to reduce redundancy in statistical testing.

It is important to note that, under certain conditions, structural input files require manual conversion before being loaded into evo3D. This includes, for example, mmCIF files generated by AlphaFold3 and structures containing non-canonical amino acids that require preprocessing. These constraints primarily reflect file-format conventions and input assumptions rather than conceptual limitations of the structural windowing framework; detailed handling strategies are described in the Methods (1.14). Finally, evo3D operates on coding-sequence MSAs and does not perform intron-exon splicing, and nucleotide alignments are therefore assumed to represent in-frame coding regions.

## Results and Discussion

We demonstrate `evo3D`’s capabilities with 3D evolutionary analysis on two protein complexes, using a set of fixed criteria for surface residue filtering, residue distance definitions, and fixed-size spatial windows (`sasa_cutoff ≥ 10`, `distance_method = “all“`, `max_patch = 15`; see Methods: Input Parameters). While adjustable by users in `run_evo3d()`, these parameters were fixed here for clarity.

### Hepatitis C Virus E1/E2 complex

The Hepatitis C virus (HCV) E1/E2 glycoprotein complex is a leading vaccine candidate, yet its genetic diversity complicates broad-coverage vaccine design (Bailey et al. 2019; Metcalf et al. 2023). To address this challenge, we applied `evo3D` as an epitope-level diversity scan, with the goal of uncovering conserved regions of an average antibody binding site (15 residues) (Reis et al. 2022). Additionally, several functional regions of E1/E2 are characterised, which allowed us to test `evo3D`’s built-in null model framework and its ability to detect these known functional regions. Finally, we repeated both analyses with linear sequence windows, allowing a direct comparison of spatial versus linear approaches.

The dataset comprised 271 HCV genotype 1b genomes obtained from BV-BRC (Olson et al. 2023), from which we extracted E1 and E2 coding regions to build two nucleotide multiple sequence alignments (MSAs). Structural context was provided by PDB ID: 8fsj (Metcalf et al. 2023). Residue numbering discussed in the text also refers to this structure file. As a preliminary dataset validation step, we calculated amino acid Shannon entropy per codon (see methods 1.11), which confirmed the expected high diversity of variable region 2 (VR2). When mapped onto structure, these values also reproduced the known general split between conserved and diverse faces of E1/E2, consistent with previous reports (Torrents de la Peña et al. 2022; Metcalf et al. 2023) (Fig. 2a).

**Figure 2.**
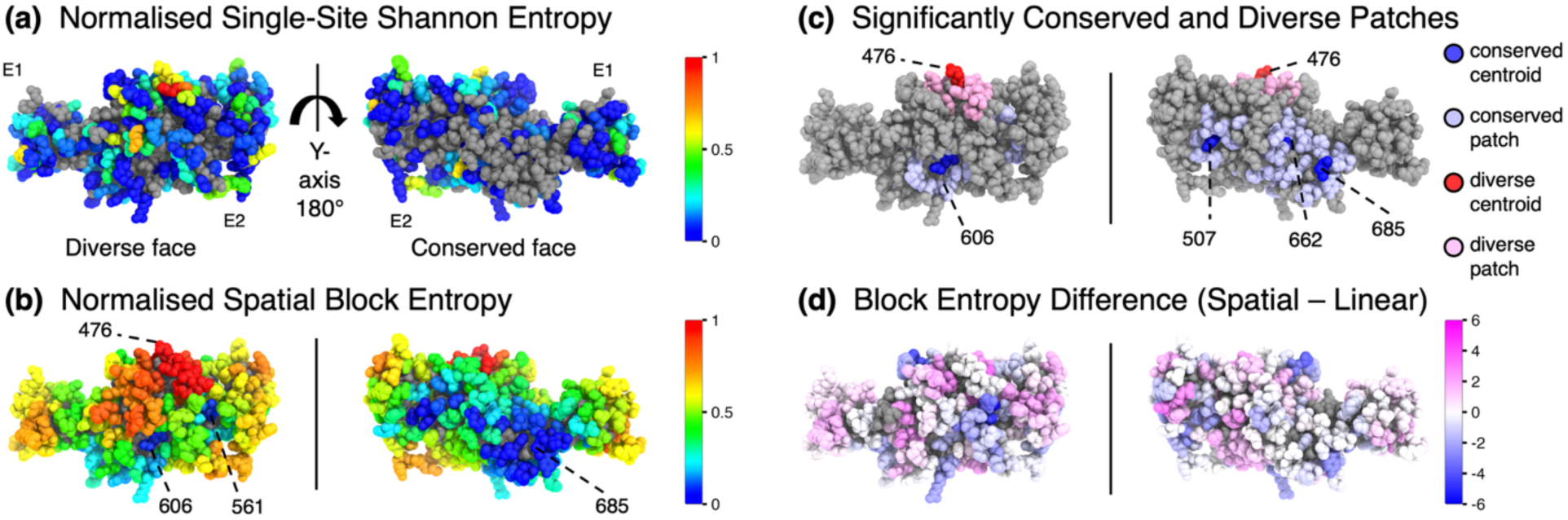
Spatial block entropy analysis of the Hepatitis C virus E1E2 complex. **(a)** Normalised single-site Shannon entropy of amino acid frequencies across 271 genotype 1b genomes, mapped onto PDB ID: 8fsj. Values are scaled between 0 and 1 by dividing by the maximum observed value of 3.4. Invariant sites are shown in grey. **(b)** Normalised spatial block entropy, calculated as Shannon entropy of haplotype frequencies for each spatial neighbourhood and then scaled between 0 and 1 by dividing by the maximum observed entropy of 7.7. Spatial neighbourhood is defined as a surface-accessible centroid residue and its 14 nearest surface-accessible neighbouring residues. The block entropy values of the neighbourhoods are mapped to centroid residues. Grey indicates buried residues excluded by the solvent-accessibility filter. Conserved neighbourhoods (E2 residues 561, 606, and 685) and a diverse neighbourhood (E2 residue 476) are annotated. Residue numbering follows PDB ID: 8fsj **(c)** Significantly conserved and diverse spatial neighbourhoods identified by comparison to an empirical null model of randomised surface haplotypes (see Methods 2.5). Four conserved patches (507, 606, 662, 685) and one diverse patch (476) were detected on E2. **(d)** Block entropy differences between spatial and linear haplotypes. Both haplotype definitions are centred on the same amino acid: linear haplotypes include ±7 residues along the sequence, while spatial haplotypes include the 14 nearest surface residues in 3D. Colours represent spatial minus linear values on a symmetric scale (–6 to +6). The maximum observed positive difference was +3.9 bits, and the maximum observed negative difference was −5.8 bits.

We next carried out an epitope-level diversity scan of the E1/E2 surface, which requires measuring the combined diversity of entire spatial neighbourhoods rather than single positions. For this, we calculated block entropy, which is defined as the Shannon entropy of amino acid haplotype frequencies (see methods 1.12). The approach has been systematically applied in T-cell vaccine research (Olsen et al. 2011; Hu et al. 2013; Raman et al. 2020; Tharanga et al. 2025) and here extended into the spatial domain of B-cell antibodies (Fig. 2b). Importantly, we identified highly conserved spatial regions on the otherwise diverse and immune-exposed face of E1/E2 (Torrents de la Peña et al. 2022), most prominently around E2 606 (block entropy = 0.6 bits) and E2 561 (0.7) (Fig. 2b). Across the E1/E2 surface, block entropy varied from complete conservation around E2 685 (0.0) to extreme diversity around E2 476 (7.7), with a median score of 3.7 bits (Fig. 2b). These results underscore the difficulty in developing a broad coverage HCV vaccination, as the median score of 3.7 bits corresponds to effectively 13 equal-frequency haplotypes sharing a typical epitope-sized region on the E1/E2 surface. However, `evo3D` identified two promising regions around E2 606 and E2 561, which are accessible to antibodies and dominated by single haplotypes covering more than 90% of the genomes sampled.

In addition to calculating spatial diversity, `evo3D` supports significance testing to identify outlier spatial neighbourhoods by generating empirical null distributions of statistics calculated from synthetic haplotypes composed of matching window sizes of resampled codons. We repurposed the block entropy scores to address two additional questions: do statistically significant regions correspond to known or inferred functions, and could they have been detected in a linear sliding window approach, in which haplotypes are defined by sequential rather than spatial proximity using windows of equivalent size (15 amino acids).

Comparing spatial block entropy scores against the null distribution revealed five significant neighbourhoods (see Methods 2.5), four conserved and one diverse (Fig. 2c). In contrast, an equivalent linear window analysis revealed only two significant regions, one conserved and one diverse, each also detected in spatial analysis. Among the significant spatial regions missed in linear analysis were two conserved neighbourhoods around E2 685 and E2 662, both of which participate in the protein-protein interface of the higher-order dimer of E1/E2 heterodimers (Fig. S1a). Additionally, the conserved region around E2 606, highlighted above as a potential broad-coverage vaccine target, was significant in spatial analysis but not in linear analysis.

Regions detected by both spatial and linear window analyses were E2 476, corresponding to the diverse VR2 segment implicated in modulating antibody recognition (Alhammad et al. 2015) and the conserved neighbourhood of E2 507, which has been suggested to contribute to E1/E2 oligomerization (Augestad et al. 2024). A complementary role for the spatial region around E2 507 may be participation in higher-order trimer interfaces, consistent with an AlphaFold3 (Abramson et al. 2024) predicted trimer (Fig. S1b) and supported by evidence that E1 forms trimers (Falson et al. 2015).

When statistical thresholds were relaxed (see Methods 2.5), the linear window analysis recovered E2 685 and E2 662 neighbourhoods, but with a reduced level of confidence compared to spatial window analysis (Benjamin-Hochberg: q = 0.044 for linear vs. q = 0.003 for spatial). However, the conserved region of E2 606 was never detected by the linear window analysis. These comparisons demonstrate that `evo3D`’s framework can detect functional regions, including conserved protein-protein interfaces and diverse immune-modulating segments. Critically, spatially-organized features were detected with greater sensitivity under spatial analysis, with the E2 606 region identified exclusively by spatial window analysis. This supports earlier findings that spatially informed methods improve the detection of 3D features (Landgraf et al. 2001; Suzuki 2004; Berglund et al. 2005).

Although spatial window analysis captured features missed by the linear window analysis, the two approaches did partially overlap in findings. Direct comparison of block entropy scores for spatial and linear windows centred on the same codon showed moderate correlation (Pearson r = 0.66), reflecting that spatial and linear windows shared an average of 7.5 out of 15 residues. However, individual windows differed substantially, sharing as few as two and as many as thirteen residues, and spatial minus linear block entropy scores ranged from −5.8 to +3.9 bits (Fig. 2d). In terms of effective haplotype diversity, this corresponds to regions nearly 60 times more conserved to 15 times more diverse in the spatial view, re-emphasizing that linear and spatial approaches capture different signals.

When the features of interest are inherently 3D, as many protein features are, the spatial perspective is not merely complementary but essential. More broadly, the HCV use cases, which included epitope-level scanning and spatial significance testing, identified conserved vaccine targets and recovered known functional features that linear window approaches overlooked. Applying `evo3D` to other protein systems requires only sequence alignment data and a structural model. and is not limited to epitope-sized windows or the block entropy statistic, making it a general-purpose framework for evolutionary analysis of 3D features.

### Chikungunya Virus E1/E2 complex

To illustrate `evo3D`’s scalability and to demonstrate several features specific to multimeric systems, we applied the workflow to the Chikungunya virus (ChikV) E1/E2 octameric complex, which is nearly an order of magnitude larger than the HCV example. This system provides a setting to highlight distinctions between window modes and analysis modes and to demonstrate how interfaces can be analysed as independent spatial haplotypes.

The spatial window analysis revealed elevated diversity adjacent to the receptor binding region of E2, but a largely conserved receptor binding region itself. This pattern is consistent with immune pressure acting on surrounding residues while functional constraints maintain conservation within the binding interface. The increased structural size produced a corresponding rise in runtime: the HCV analysis completed in five seconds, whereas the ChikV analysis required approximately 1.5 minutes. Most of this increase was due to the construction of the all-atom distance matrix. When distances were computed using Cα or centroid coordinates, runtime decreased to approximately 20 seconds.

We obtained 379 ChikV genomes from BV-BRC (Olson et al. 2023), constructed nucleotide MSAs for E1 and E2, and used the octameric structure PDB ID: 8fcg (Chmielewski et al. 2024) for structural context and residue referencing in the text. Each MSA mapped to four chain copies (Fig. 3a), producing four distinct spatial environments per codon. This system illustrates the window and analysis-mode distinctions formalised in `evo3D`: whether spatial windows retain all residues or are deduplicated to unique codons (window mode), and whether analysis assigns one spatial window for every residue or one for every codon (analysis mode). Because block entropy is unaffected by repeated codons, we constructed residue-mode windows containing 15 residues. Only 1.2 percent of windows contained codon duplicates, with some windows comprising as few as ten unique codons among fifteen residues. Although infrequent, such cases would distort any spatially averaged statistic, highlighting the need for explicit control over window mode.

**Figure 3.**
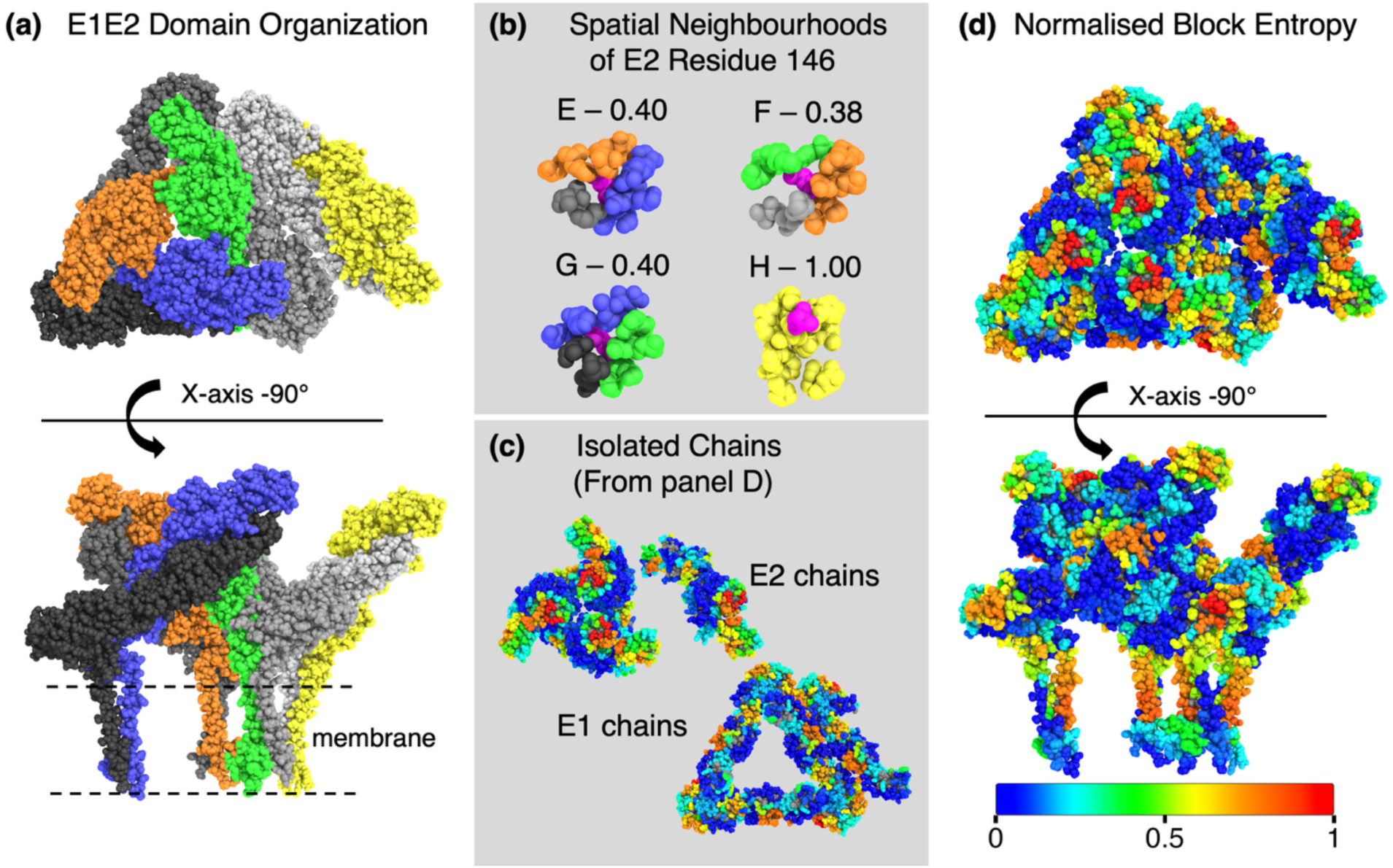
evo3D analysis of large multimeric complexes illustrated with the Chikungunya virus E1E2 octamer. **(a)** Domain organization of the E1E2 octamer, composed of four E1E2 heterodimers. E2 chains are shown in blue, orange, green, and yellow; E1 chains are shown in shades of grey. **(b)** Spatial neighbourhoods defined for residue 146 of E2 (8fcg numbering), with residue 146 in magenta. Chain colours are consistent with panel A. The source chain of residue 146 and the block entropy of each neighbourhood are indicated above the corresponding window. Because each codon is represented in four chains, values in (c) and (d) are merged across chains, such that the displayed block entropy was averaged across the four neighbourhoods. **(c)** Isolated E1 and E2 chains from (d), shown separately for clarity. Spatial neighbourhoods can cross E1 and E2 chains, as illustrated in panel B. **(d)** Normalized block entropy, values are scaled to the observed maximum of 2.2. Grey residues in (c) and (d) are solvent-inaccessible and therefore excluded from spatial neighbourhoods.

To evaluate how consistently the same codon is represented across its four structural environments, we performed the analysis in residue mode, assigning a separate window and score to every residue-level environment and therefore generating four scores per codon (Fig. 3b). Comparing the maximum and minimum block entropy values for each codon showed that the four environments were generally consistent (median difference = 0.03 bits, IQR = 0–0.12). Fourteen codons, however, showed differences of at least 1.0 bit, with a maximum difference of 1.7 bits. These larger differences reflect chain-specific variation in solvent exposure of the neighbouring residues. The octamer structure contains a rich set of burial and exposure contexts each codon encounters in the intact viral envelope, but these contexts are distributed across different chains. By switching to codon analysis mode, `evo3D` can merge these chain-specific environments into a single codon-level representation that captures the complete structural context for each codon (see Methods 1.8).

After comparing residue-level environments, we collapsed these values into a single codon-level score by averaging the block entropy values across chains (Fig. 3c-d). Overall diversity ranged from 0 to 2.2 bits (median 0.4), with E2 showing higher diversity (median 0.6) than E1 (median 0.2). The highest-diversity window was centred on E2 residue 119, which is notable because residue 119 forms a contact with MXRA8, the human entry receptor (Zhang et al. 2018; Song et al. 2019), as determined from PDB ID: 6jo8 (Fig. S2a). Although this window represents the most diverse local environment on the E1/E2 surface, E2 residue 119 itself is completely invariant. This illustrates a key property of spatial windows: block entropy measures the diversity of the three-dimensional neighbourhood, rather than the central codon anchoring the window.

To characterize the MXRA8 interface directly, `evo3D` supports constructing spatial haplotypes for specified interfaces. Interfaces were included by setting `interface_chain = c(“M”,“N”,“O”)` in the wrapper call. Using PDB ID: 6jo8 (Song et al. 2019), where chains M, N, and O correspond to MXRA8, contacts within 4 Å of MXRA8 and E1/E2 were extracted. Because block entropy is window-size dependent, we compared per-site amino acid Shannon entropy of MXRA8-contacting codons to that of the full E1/E2 surface (Fig. S2b). Interface codons were strongly conserved: only one exceeded 1 bit (E2 residue 74, 1.2 bits), while four codons exceeded 1 bit on the full E1/E2 surface (maximum at E1 residue 211, 1.6 bits). The interface had a median per-site entropy of 0 and an interquartile range of 0 to 0.03, slightly higher than the surface as a whole (median 0; IQR 0–0). This highlights the importance of treating interfaces as their own analytical units. The highest-diversity spatial window on E1/E2 is centred on a residue that contacts MXRA8, but that residue is invariant and the MXRA8-binding site is largely invariant. Thus, diversity is highest near the MXRA8-binding surface, not within it.

This example illustrates the scalability of `evo3D` to large protein complexes. The wrapper supports diverse structural contexts, provides access to multiple window and analysis modes, and supports interfaces as separate analytical units. Applied to the ChikV octamer, these components allow chain-specific environments to be compared directly or merged into unified codon-level contexts and allow the MXRA8-binding interface to be examined as its own spatial haplotype. The full analysis, which mapped two MSAs to four chains each and constructed spatial haplotypes for every surface residue, can be completed in approximately 20 seconds on a laptop with an Intel Core i7-13620H CPU with roughly 1 GB memory usage (see Supplementary Methods S.6 and Table S4) `evo3D` extends structure-informed analysis to multimeric assemblies and interface-level contexts while remaining computationally tractable on standard hardware.

Although the two application examples focus on intraspecific comparisons of viral proteins, evo3D is not limited to viral proteins or to within-species MSAs. Between-species analyses, such as detecting conservation of enzymatic domains or protein-protein interfaces, or detecting per-codon signals within spatially coherent regions, remain feasible because protein structure often diverges more slowly than sequence similarity (Rost 1997; Illergård et al. 2009). The practical boundary of such analyses is the variability within the MSA, particularly the frequency of insertions and deletions. When indels are prevalent, spatial windows derived from a single structural model may no longer accurately represent the structural context across the full alignment.

## Conclusion

`evo3D` provides a practical framework for linking MSAs to three-dimensional structure and returns spatially defined haplotypes for downstream statistics. The package is designed to support a wide range of applications, from basic evolutionary summaries to more specialized analyses of immune recognition, viral diversification, and scales from monomers to multimer assemblies. By formalizing the structural integration layer and exposing spatial haplotypes for downstream analysis, evo3D makes structure-informed evolutionary analysis both tractable and reproducible across diverse protein systems.

## Methods

### 1. Package details

#### 1.1 Workflow overview

`evo3D` consists of a set of modular components that perform each step of structure-informed evolutionary analysis. These modules are described in the Implementation section, but in practice, they are orchestrated through the `run_evo3d()` wrapper, which removes technical barriers and provides a unified interface. The wrapper accepts one or many MSA and PDB inputs, builds an internal run grid that maps PDB chains to MSAs through fast k-mer similarity, and executes the full analysis in a single, seamless workflow (Fig. 1).

#### 1.2 Parameter-control structure

The behaviour of `evo3D`’s underlying modules is governed by named control lists supplied to `run_evo3d()`. Controls are grouped according to the stage of the workflow they modify: `msa_controls`, `pdb_controls`, `aln_controls`, `collapse_controls`, `stat_controls`, and `output_controls`. Each list contains only the parameters relevant to its corresponding modules, allowing the wrapper to remain compact while supporting extensive customization. Default values for all controls can be viewed with `show_evo3d_defaults()`, and individual settings may be overridden as needed.

#### 1.3 Automatic chain mapping

Chain assignment is performed only when the `run_evo3d()` chain argument is set to `“auto“` (the default). In this mode, `run_evo3d()` identifies which structural chain corresponds to each MSA during run-grid construction. Matching is based on fast k-mer similarity between the MSA-derived peptide reference and the amino-acid sequences of each chain in the structural model; chains are ranked by k-mer overlap, and the highest-scoring chain is selected. These assignments are stored explicitly within each run-grid entry and are used by all downstream modules. Users may bypass automatic mapping entirely by supplying chain specifications directly through the chain argument (see Supplementary Methods S.1.1).

#### 1.4 MSA processing and reference construction (msa_to_ref)

The `msa_to_ref()` module standardizes user-supplied MSAs and generates the peptide reference sequence used for aligning the MSA to the PDB. Inputs may be nucleotide or amino-acid MSAs provided as matrices (for example, from `bio3d::read.fasta()`) or as file paths. The module detects sequence type automatically by examining the most frequent character types for the first 100 columns of input MSA.

A reference sequence is then constructed according to parameters defined in `msa_controls`. Users may select the least-gapped sequence in the alignment, a specified MSA row, or a consensus sequence computed from non-gap characters. Nucleotide MSAs are translated to amino acids using a user-specified genetic code. The module returns a standardized S3 object containing the original MSA, the reference sequence, and the translated reference peptide. These outputs form the sequence foundation for all subsequent MSA to PDB alignment steps.

#### 1.5 Structural parsing and spatial-window generation (pdb_to_patch)

The `pdb_to_patch()` module parses structural files and constructs residue indexed spatial windows that define the three-dimensional neighbourhoods used throughout `evo3D`. Inputs may be PDB or mmCIF file paths or pre-loaded objects from `bio3d::read.pdb()` or `bio3d::read.cif()`. The module first validates file format and standardizes fields such as insertion codes to ensure consistent residue indexing across chains.

Sequences from the user or wrapper specified chains are extracted, and inter-residue distances are computed according to the method specified in `pdb_controls`. Users may select minimum all-atom distances, Cα–Cα distances, or distances between residue centroids. Solvent accessibility is then calculated with the DSSP-style algorithm (see section 1.6) incorporated into `evo3D`, and residues are classified as exposed or buried based on user-defined thresholds. Optional identification of inter-chain contacts, guided by the `interface_chain` argument, allows interface-specific windows to be generated when required.

Using these distance and exposure measures, the module constructs spatial windows around each residue according to either fixed size or distance cutoff rules. These windows are defined entirely at the structural level, before any reference to the MSA, and therefore reflect the geometric properties of the input model. The output is a standardized S3 object containing the structural data, the chain sequences, distance matrices, and a table of residue centred spatial windows and exposure values for each residue.

#### 1.6 Solvent-accessibility calculation

Solvent-accessible surface area (SASA) was computed with a DSSP-style algorithm re-implemented in C++ and integrated into `evo3D` through the function `.calculate_accessibility()`. The implementation reproduces the geometric definitions of DSSP (Kabsch and Sander 1983), including atom-specific van der Waals radii and geodesic-sphere sampling of atomic surfaces to estimate accessibility. Validation against MKDSSP v4.4.10 (Hekkelman et al. 2025) across 95 protein structures (114,140 residues) showed near-perfect agreement (Pearson r = 0.9999784) with per-residue deviations of −0.5 to +0.5 Å², well within the ≈1 Å² numerical variance reported in the original DSSP publication (Kabsch and Sander 1983).

#### 1.7 MSA-to-structure alignment and codon mapping (aln_msa_to_pdb)

The `aln_msa_to_pdb()` module aligns the reference peptide generated by `msa_to_ref()` to the residue sequence extracted from each structural chain and establishes the mapping between MSA codons and PDB residues. Alignment is performed with the `msa` package (Bodenhofer et al. 2015), which provides the Clustal Omega algorithm (Sievers et al. 2011). The resulting alignment table is stored and returned to the user at the end of the workflow. If a user modifies this alignment, `run_evo3d()` can be restarted with the revised table, which provides a fully transparent and reproducible mapping process (see Supplementary Methods S.1.3 and Table S1).

The residue-indexed windows produced by `pdb_to_patch()` are then converted into codon-indexed windows. Each residue is assigned a codon position based on the reference alignment. Residues that align to gaps in the MSA reference do not have corresponding codons and are removed from codon-indexed windows. Codon duplicates that arise in multimeric assemblies are retained or removed according to the deduplication rules specified in `aln_controls`. When fixed-count windows are requested, the module reconstructs any windows that fall below the target size so that downstream analyses operate on consistent codon counts.

The module returns a standardized S3 object that includes a table of codon-to-residue mappings and codon indexed windows for all residues in the specified PDB chains, the individual MSA to PDB chain alignments for optional user correction, and a coverage list summarizing the completeness and identity of each MSA relative to each structural chain.

#### 1.8 Codon-level window merging and multimer integration (collapse_to_codon)

The `collapse_to_codon()` module converts the residue-level table produced by `aln_msa_to_pdb()` into a codon-level table. In multimeric assemblies or multi-model analyses, a single codon may appear multiple times because it maps to several structural residues. For example, the same codon may correspond to positions 127 in chains E, F, and G, each with its own residue-centred window. `collapse_to_codon()` condenses these separate neighbourhoods into one representation per codon and merges their windows according to the mode specified in `collapse_controls`. The available modes are `union`, `distance`, and `exposure_distance`. Detailed definitions of each mode are provided in the Supplementary Methods (S.2).

#### 1.9 Extraction of spatial haplotypes (extract_msa_subsets)

The `extract_msa_subsets()` is a submodular function generally applied with the `aln_msa_to_pdb()` or `collapse_to_codon()` module, but within the wrapper function `run_evo3d(),` it is placed after the `collapse_to_codon()` module. This function extracts the MSA subsets corresponding to each codon-indexed window and returns them as spatial haplotypes. For every window, the module gathers the codon positions defined by `aln_msa_to_pdb()` and `collapse_to_codon()` and slices the original MSAs to produce discontinuous but structurally coherent subsets that serve as inputs for downstream statistics.

Haplotype construction across multiple MSAs is controlled by the `use_sample_names` parameter in `aln_controls`. When this option is set to `TRUE`, discontinuous sequence segments are concatenated only where row names are identical. Sequences from one MSA with no identical name in other MSAs are dropped. This ensures that codons drawn from different genes or loci originate from the same genome. When `use_sample_names` is `FALSE`, MSA subsets are concatenated row-wise, and any sequences left unpaired after exhausting one MSA are padded with gap characters.

#### 1.10 Calculation of window-level statistics (calculate_patch_stats)

The `calculate_patch_stats()` module provides six built-in statistics that can be applied directly within the `evo3D` workflow. These include Shannon entropy for single sites, averaging single-site Shannon entropy per patch, and block entropy, along with several summary statistics from the `pegas` (Paradis 2010) package, such as nucleotide diversity, Tajima’s *D*, and haplotype diversity. These functions are intended as convenient defaults rather than a complete analysis suite.

Because `evo3D` returns all spatial haplotypes to the user, most statistical analyses can be applied independently outside the wrapper. Users may compute custom metric directly on the returned spatial haplotypes without modifying the `evo3D` workflow. The internal statistics module, therefore, serves primarily as an optional shortcut for standard diversity measures.

#### 1.11 Single-site entropy (Fig. 2a, Fig. S2)

Multiple sequence alignments were translated to amino acid sequences using the R package `seqinr` (Charif and Lobry 2007), with translation tables specified by the `genetic_code` parameter in `msa_controls` when alternative codon systems were required. Shannon entropy (Shannon 1948) was calculated for each alignment position from observed amino-acid frequencies. Gaps and ambiguous residues (“X”) were excluded on a per-position basis.

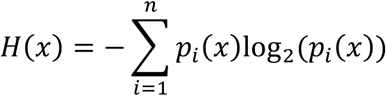

Where *H* is Shannon entropy, *x* denotes the alignment position, *i* represents an amino acid type, *n* is the number of distinct amino acid residues at position x, and *p_i_(x)* is the observed frequency of amino acid *i* at position *x*.

#### 1.12 Block entropy (Fig. 2b, Fig. 3b-d)

As described in Olsen *et al*. (Olsen et al. 2011), haplotype-level Shannon entropy (block entropy) was computed from the frequency distribution of amino acid haplotypes within spatial or linear windows. Haplotypes were translated to amino acids using seqinr, and sequences containing gaps or ambiguous residues were excluded before frequency estimation.

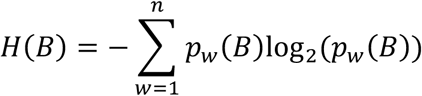

Where *H* is Shannon entropy, *B* denotes the MSA subset, *w* represents a haplotype, *n* is the number of distinct haplotypes in MSA subset *B*, and *p_w_(B)* is the observed frequency of the haplotype in the MSA subset *B*.

#### 1.13 Return structure of run_evo3d()

The `run_evo3d()` wrapper returns a structured S3 list that contains all sequence, structure, alignment, and window-level information produced during the analysis. This list serves as both the final output and a complete record of all intermediate steps, which allows users to inspect components, rerun modules with corrected inputs, or perform additional analyses directly on the returned objects.

The return object includes:

- `evo3d_df` A table of the final MSA-PDB alignment, codon-indexed windows, and optional statistics per window.
- `final_msa_subsets` All spatial haplotypes extracted for each window.
- `msa_info_sets` output objects from runs of `msa_to_ref()`
- `pdb_info_sets` output objects from runs of `pdb_to_patch()`
- `aln_info_sets` output objects from runs of `aln_msa_to_pdb()`
- `call_info` A full record of parameters supplied to `run_evo3d()`, including all control settings used during the analysis, and the run grid built internally.

This return structure provides full transparency across all stages of the workflow and gives users direct access to both spatial haplotypes and the intermediate tables needed for independent or extended analyses.

#### 1.14 File Preprocessing

Certain types of structural input files require manual conversion before being loaded into `evo3D`. Currently, the mmCIF reader in `bio3d` has incomplete support for recent formats produced by AlphaFold3, which may cause some structures to be parsed incorrectly. Converting AlphaFold3 mmCIF outputs to PDB format using tools such as GEMMI (Wojdyr 2022) resolves this issue. Future versions of `evo3D` plan to implement a dedicated parser that removes this user intervention. Second, non-canonical amino acids present as HETATM records in structural files will not be incorporated into sequence parsing or solvent-accessibility calculations in evo3D. These residues can be converted to ATOM records prior to analysis to be included in sequence and window generation. When converted to ATOM, these residues will appear as “X” in the MSA-to-structure mapping and will be present in downstream windows, corresponding to valid codon columns in the MSA. A more complete solution would assign a non-canonical residue to the closest canonical type, but this requires user decisions on how to convert non-canonical residues. Finally, the workflow assumes coding-strand MSAs and does not handle exon-intron splicing. Users working with genomic alignments should preprocess their data to generate coding-only MSAs before supplying them to `evo3D`.

### 2. Analysis details

#### 2.1 Hepatitis C Virus (HCV) genetic data

HCV genotype 1b genomes were retrieved from BV-BRC (accessed July 24 2025; n = 274; (Olson et al. 2023)) and formatted as a local nucleotide BLAST database. A representative SwissProt genotype 1b polyprotein (accession O92972; (The UniProt Consortium 2025)) was trimmed to the E1-E2 region and used as the tblastn query (BLAST+ v2.13.0; (Camacho et al. 2009)). Hits spanning the full-length query were retained. One partial hit beginning at query position two was manually extended by one codon to recover the complete region, yielding 273 full-length matches. Corresponding nucleotide segments were extracted with blastdbcmd and translated with SeqKit (v2.3.1; (Shen et al. 2024)). Sequences containing internal stop codons or fewer than 95% standard amino acid characters were removed, leaving 271 genomes. The E1-E2 amino acid reference sequence (O92972) was appended before multiple sequence alignment (MSA) with MUSCLE (v5.1; (Edgar 2022)). The amino acid alignment was mapped to nucleotide sequences to insert triplet gaps and preserve the reading frame. Alignment columns with >90% gaps were removed. Based on SwissProt-defined gene boundaries, the final E1 and E2 nucleotide MSAs were extracted for downstream analysis.

#### 2.2 Chikungunya virus (ChikV) genetic data

ChikV genomes were retrieved from BV-BRC (accessed October 30, 2025; n = 1,139; (Olson et al. 2023)) and filtered to retain sequences collected between 2016 and 2025 (n = 632). From these, up to 50 genomes per sampling year were selected, yielding 385 genomes, which were formatted as a local nucleotide BLAST database. A representative SwissProt ChikV polyprotein (accession Q5XXP3; (The UniProt Consortium 2025)) was trimmed to the E2-E1 region and used as the tblastn query (BLAST+ v2.13.0; (Camacho et al. 2009)). Hits spanning the full-length query were retained, producing 380 complete matches. The corresponding nucleotide regions were extracted with blastdbcmd and translated with SeqKit (v2.3.1; (Shen et al. 2024)). Sequences containing internal stop codons or fewer than 95% standard amino acid characters were removed, leaving 379 genomes. The E2-E1 amino acid reference sequence (Q5XXP3) was appended before multiple sequence alignment with MUSCLE (v5.1; (Edgar 2022)). The amino acid alignment was mapped to nucleotide sequences to insert triplet gaps and preserve the reading frame. Alignment columns with >90% gaps were removed. Based on SwissProt-defined gene boundaries, the final E1 and E2 nucleotide MSAs were extracted for downstream analysis.

#### 2.3 Parameterization for HCV and ChikV examples

HCV and ChikV analyses were performed using the run_`evo3d`() wrapper with the following parameters specified in the argument `pdb_controls = list(max_patch = 15, distance_method = “all”, and sasa_cutoff = 10)`. The `max_patch` parameter defines fixed-size spatial windows containing the specified number of residues. The `distance_method` parameter determines how inter-residue distances are calculated: `“all“` uses the minimum distance between all non-hydrogen atom pairs. The `sasa_cutoff` parameter filters residues by solvent accessibility, retaining only those with solvent-accessible surface area (SASA) above the defined threshold.

#### 2.4 Linear sliding windows (HCV)

To ensure a direct comparison between spatial and linear window analyses, identically sized (15-amino acid) linear sliding windows were generated across each MSA. Window centres were restricted to the start and end of resolved regions in the structural model PDB ID: 8fsj (Metcalf et al. 2023), preventing inclusion of regions present in the MSA but outside the resolved region of the structural model, such as transmembrane domains and the highly diverse HVR1 segment. At the boundaries of the resolved region, linear windows were permitted to extend into adjacent MSA positions that lacked structural coordinates. Thus, spatial and linear analysis were matched in window size, centre residue, and regional constraints, differing only in how residue window inclusion was defined.

#### 2.5 Null-model generation and significance testing (Fig. 2c)

To test for significance in conserved and diverse windows, we generated null distributions of block entropy statistics using synthetic haplotypes. Following Berglund *et al*. (2005), one of the earliest studies to formalise spatial sliding-window analysis, we pruned overlapping windows to reduce redundancy arising from highly similar spatial neighborhoods and reported significance under two tiers of p-value correction for multiple testing. The null distributions were generated using the same procedure for spatial and linear window analyses, including the spatial window analysis of the HCV E1/E2 complex and the linear window analyses of E1 and E2 separately.

Using the `generate_null_model()` function in `evo3D`, 10,000 synthetic windows of 15 codon positions were generated, matching the window size used in the corresponding analysis. Codon positions available for sampling were those present in analysis windows. Codon positions were sampled uniformly from this set, and the resulting windows were used to extract synthetic haplotypes from the analysis MSA(s). Block entropy was calculated for each synthetic haplotype to generate null distributions. For each analysis window, a two-sided empirical p-value was calculated from the null distribution using the proportion of simulated block entropy values at least as extreme as the observed value, with a +1 pseudocount applied to the numerator and denominator to avoid p-values of zero (Phipson and Smyth 2010). Standardized effect sizes were calculated as Z-scores relative to the null mean and standard deviation.

To prune overlapping windows with high similarity, windows were first ranked by absolute block entropy Z-score (|z|), and overlapping windows (≥33% shared codons) were iteratively pruned in descending order of |z| using the function `filter_overlaps()`. Berglund *et al*. (2005) explored overlap thresholds of 10%, 33%, and 80%. We adopted the intermediate threshold of 33% as a balance between stringent and permissive pruning. Empirical p-values were adjusted using the Benjamini-Hochberg (BH, α = 0.05) and Bonferroni (α = 0.1) corrections (Benjamini and Hochberg 1995). Two significance tiers were defined: a stringent threshold requiring significance under both corrections and a relaxed threshold requiring significance under BH only.

#### 2.6 Comparison of spatial and linear entropy (Fig. 2d)

Differences between spatial and linear block entropy were calculated as (spatial minus linear) per window, enabling direct comparison of the two window approaches centred on the same amino acid.

#### 2.7 Multimeric averaging and codon-collapsing modes (Fig. 3c-d)

For the ChikV multimer, entropy values from residue-centred windows were averaged across all residues mapped to each codon, producing a single representative score per codon. This approach merges values but not window contexts, as each codon still spans multiple residue-centred windows. More direct codon-collapsing methods, `union`, `distance`, and `exposure_distance`, are implemented in `evo3D` to merge residue-centred windows into unified codon-level representations.

#### 2.8 Interface capture of ChikV E1-E2 and Mxra8 (Fig. S2)

Interface residues were identified using the CHIKV E1/E2-MXRA8 complex PDB ID: 6jo8 (Song et al. 2019). Inter-chain contacts were defined by a 4 Å distance threshold (`interface_dist_cutoff,` default = 4) specified in `pdb_controls`.

## Acknowledgments

The authors thank Ilinca Ciubotariu and Giovanna Carpi for an earlier collaboration that shaped our appreciation for structure-informed analysis and motivated this work. We also acknowledge the support of the Rosen Center for Advanced Computing at Purdue University. Last, we thank the reviewers for their suggestions that improved the manuscript and better positioned evo3D into the broader structure-informed methodological landscape.

## Author Contributions

B.K.B.: Conceptualisation, Data curation, Formal analysis, Software, Writing; Q.H.: Conceptualisation, Funding acquisition, Supervision, Writing

## Data Availability

`evo3D` package code is available at https://github.com/bbroyle/evo3D, and can be installed through `devtools::install_github(‘bbroyle/evo3D’)`. Data and code for all results and supplementary results are available at https://zenodo.org/records/18986474 (doi: 10.5281/zenodo.18986474).

## Supplementary Figure Legends

**Figure S1. Accessibility of significantly conserved and diverse surface patches in the Hepatitis C virus E1E2 complex. (a)** Significantly conserved (blue) and diverse (red) neighbourhoods identified in Fig. 2c (PDB ID: 8fsj) are shown superimposed on the experimentally solved E1E2 dimer (PDB ID: 8rjj; RMSD = 1.295 across 2043 atoms). The neighbourhood of residue 606 is surface-accessible, while the neighbourhood of residue 507 is likely less accessible as an epitope due to steric hindrance. The neighbourhood of residue 476 is diverse and fully accessible, consistent with known frequent immune recognition. **(b)** The same significant patches mapped to the predicted trimer by superimposing 8fsj onto an AlphaFold3-predicted trimer (RMSD = 2.091 across 2176 atoms). Neighbourhoods for residues 606 and 476 are predicted to be accessible, while the conserved neighbourhood around residue 507 participates in the predicted trimer interface.

**Figure S2. Interface-aware capture in `evo3D` enables binding interfaces to be analysed as spatial neighbourhoods. (a)** MXRA8 (pink) bound to the Chikungunya virus E1/E2 complex (PDB ID: 6jo8), with the viral surface coloured by normalised block-entropy values derived from spatial windows constructed on the octameric assembly (PDB ID: 8fcg) (see Fig. 3d). This visualisation shows how variable and conserved spatial patches identified in Fig. 3d relate to the MXRA8-binding footprint. The grey region in the upper left corresponds to the E3 protein, for which no MSA information was generated in this study. **(b)** Per-site amino acid entropy across the three copies of E1/E2 - MXRA8 binding interface compared to per-site entropy of the full E1/E2 surface (computed using 8fcg). Interface contacts were defined from 6jo8. Because both 6jo8 and 8fcg map to the same MSAs, codon positions are directly comparable, allowing an interface defined in one structure to be evaluated on the other.

