## Supplementary Information for "evo3D R package: a spatial haplotype framework for structure-informed analysis of molecular evolution"

This PDF file includes:

Supplementary Methods

Figs. S1 to S2

Tables S1 to S4

SI References

Supplementary Methods
**S.1: evo3D wrapper: run_evo3d() input control and parameter space**
run_evo3d() is the top-level wrapper that links sequence and structure, builds codon-aligned, structure-aware windows, subsets multiple sequence alignments (MSAs) to form spatial haplotypes, and optionally computes diversity and neutrality statistics. Internally, it orchestrates the MSA, PDB, alignment, codon-collapse, and statistics modules in a single end-to-end pipeline. The wrapper is designed to minimise technical barriers and to handle most analyses automatically; however, manual control of the MSA-to-PDB chain mappings and of many tunable parameters are available to users.

**S.1.1: chain, interface_chain, occlusion_chain manual control**
PDB chains corresponding to each input multiple sequence alignment (MSA) are resolved automatically by default, but manual control is available via the chain argument. Additional chain-related arguments (interface_chain and occlusion_chain) are not automatically detected; they define interfaces as their own spatial windows and solvent-accessibility-modifying chains, respectively. An example run with manual control of these three chain parameters follows:

```
run_evo3d(
	msa = list(msa1, msa2),
	pdb = list(pdb1, pdb2),
	chain = list(c(‘auto’, ‘BC’), c(‘AB’, NA)),
	interface_chain = list(c(‘EF’, ‘G’)),
	occlusion_chain = list(‘MNO’, c(‘M’, ‘N’, ‘O’))
)
```

The chain argument is supplied as a list parallel to the PDB input, with each inner vector providing one entry per MSA. Thus chain[[1]] = c("auto","BC") applies to pdb1, meaning that msa1 uses automatic chain detection while msa2 maps directly to chains B and C. Likewise, chain[[2]] = c("AB",NA) applies to pdb2, mapping msa1 to chains A and B and omitting msa2.

The occlusion_chain argument allows chains to influence the solvent accessibility of analysis chains. It is also suited to handling HETATMs as occluding entries, allowing glycans and other small molecules to modify solvent accessibility. The argument is supplied as a list where each inner vector is positionally assigned to the input PDB files. Under this convention, "MNO" and c("M","N","O") are interpreted identically, both indicating that chains M, N, and O act as occluding neighbours (as in the example, "MNO" for pdb1 and c("M","N","O") for pdb2).

The interface_chain argument is also supplied as a list where each inner vector is positionally assigned to the input PDB files; however, interface_chain uses concatenated identifiers to define combined interfaces. For example, interface_chain[[1]] = c("EF","G") instructs evo3D to build two interface windows for pdb1: one merging contacts from chains E and F into a single spatial haplotype, and a second for contacts onto chain G. Supplying c("E","F","G") instead would produce three separate interface haplotypes. The second element interface_chain[[2]] is left blank, instructing no interfaces for pdb2 and is equivalent to interface_chain[[2]] = NA. Any elements left blank for chain argument are assumed to be “auto”. Any left blank for occlusion_chain or interface_chain are assumed to be NA. Interface contacts, and therefore interface-derived spatial haplotypes, are computed only for chains that participate in the MSA mapping; chains with no MSA mapping are ignored.

This parsing logic is implemented in the internal function .setup_chain_mapping() (accessible via evo3D:::.setup_chain_mapping()). A practical implication of this design is that PDB chain identifiers must be single characters in order to be compatible with manual chain specification.

**S.1.2: Additional arguments to run_evo3d()**
Several additional arguments control the wrapper's behaviour. Extending the code example from **S.1.1**, the following call illustrates all wrapper-level options that users may adjust:

```
run_evo3d(
	…,
	analysis_mode = ‘residue’,
	calculate_stats = FALSE,
	detail_level = 2,
	verbose = 2
)
```

The analysis_mode argument defaults to "codon", which produces spatial haplotypes and a results data frame indexed at the codon level. Setting analysis_mode = "residue" instead treats residues within each PDB as the indexing unit. The conceptual and practical implications of these two modes are described in the main text and expanded in **S.2**.

The calculate_stats argument is TRUE by default and triggers the downstream statistics module. Because this module is often the longest component of the overall runtime, especially for large patch sets, setting calculate_stats = FALSE provides a useful shortcut when only structural mappings or patch definitions are required.

The detail_level argument controls the amount of information retained in the returned object. Returning the full set of patch-level MSAs can easily exceed 1 GB of memory, so reducing the level can greatly reduce memory usage. detail_level ≥ 1 is required for alignment correction and analysis restarting (see S.1.3).

- detail_level = 2 retains all spatial haplotypes (patch-level MSAs) in R, and PDB distance matrices.
- detail_level = 1 removes the final patch-level MSAs but keeps the PDB distance matrices as well as the input MSA and PDB objects.
- detail_level = 0 drops the input PDB and MSA objects and PDB distance matrices, but retains the full results table and internal MSA–PDB alignment mappings for inspection.
- detail_level = -1 returns nothing and is suitable for batch processing when outputs are written to file only.

The verbose argument controls the amount of messaging printed during execution. verbose = 2 reports module and submodule-level progress; verbose = 1 reports only module-level progress; and verbose = 0 suppresses all routine messages except errors, stops, and warnings.

**S.1.3: Restarting run_evo3d() with corrected alignments**
MSA to PDB alignments for each MSA–structure combination are stored in aln_info_sets. These objects are organised by PDB, such that aln_info_sets$pdb1$pos_mat contains the alignment of every analysis MSA to the chains of pdb1. Shifting single or row blocks is accomplished with adjust_aln()or with any spreadsheet editing software. Misalignments, when they occur, typically arise near unresolved or missing regions of the PDB structure (see Table S1).

**Table S1.** **Example of a misaligned residue**. residue_id 3_A_ should align with codon 47, but is instead aligned to codon 50. The error occurs in gappy regions of the alignment, where both gap-open and gap-end positions yield equivalent alignment scores. Corrections could be performed in R or any spreadsheet editing software.

| codon | residue_id | ref_aa | pdb_aa |
| --- | --- | --- | --- |
| 45 | 1_A_ | G | G |
| 46 | 2_A_ | S | S |
| 47* | - | K | - |
| 48 | - | S | - |
| 49 | - | P | - |
| 50* | 3_A_ | K | K |
| 51 | 7_A_ | H | H |
| 52 | 8_A_ | T | T |

*: refers to rows that need to be manually corrected

To restart an analysis after manually correcting an alignment, the user replaces the relevant aln_info_sets[[pdb_name]]$pos_mat with the corrected alignment table and supplies the entire results list (including msa_info_sets, pdb_info_sets, and results data frame) from the previous run back to the wrapper via the restart_run argument:

run_evo3d(restart_run = correction)

When restart_run is provided, run_evo3d() resumes the pipeline within the aln_msa_to_pdb() module, using the user-corrected alignments in place of the original ones. Any new arguments passed to the wrapper are ignored; the analysis proceeds with the parameters from the previous run.

**S.1.4: Full parameter space of control lists**
The run_evo3d() wrapper provides a highly flexible workflow that is tuneable by users through control lists for each module. To view a full list of arguments to these control lists, users may run: show_evo3d_defaults(), or to see a specific set show_evo3d_defaults(‘pdb’).

run_evo3d(
…,
msa_controls = list(ref_method = 1),
pdb_controls = list(max_patch = 20, sasa_cutoff = 10),
aln_controls = list(use_sample_names = FALSE),
collapse_controls = list(merge_type = ‘union’),
stat_controls = list(calc_block_entropy = TRUE),
output_controls = list(output_dir = ‘.’),
)

The parameter space is as follows (**Table S2**)

**Table S2. The complete parameter list of run_evo3d() wrapper.**

| Module | Parameter | Arguments (defaults listed first) | Usage |
| --- | --- | --- | --- |
| msa_controls | ref_method | “consensus”, “least_gap”, numeric() | construct per position non-gap consensus sequence, or use least gap sequence as reference, or use specific row as reference |
| msa_controls | force_seq_type | NULL, “nucleotide”, “protein” | To skip automatic sequence type detection |
| msa_controls | genetic_code | 1, numeric() | The ncbi genetic code for translation, this parameter is passed to seqinr::translate() (Charif and Lobry 2007) |
| pdb_controls | distance_method | “ca”, “centroid”, “all” | Determining inter-residue distances from Cα, or heavy atom centroids, or minimum distance between all heavy atoms |
| pdb_controls | dist_cutoff | 15, numeric() | Distance defining inclusion in spatial windows |
| pdb_controls | max_patch | NA, numeric() | Number or residues or codons for fixed count windows, can be provided with dist_cutoff to form fixed count windows as long as they do not exceed a defined distance |
| pdb_controls | patch_mode | “codon”, “residue” | Determines if windows deduplicate codons or retain duplicates, additionally for fixed count windows controls if it is residue or codon count. |
| pdb_controls | rsa_cutoff | 0.1, numeric(), NA | Solvent accessibility cutoff for spatial window seed residues |
| pdb_controls | sasa_cutoff | NA, numeric() | Solvent accessibility cutoff for spatial window seed residues, can be combined with rsa_cutoff |
| pdb_controls | rsa_method | “rose”, “miller”, “theoretical_tien”, “empirical_tien” | Maximum solvent accessibility values per amino acid, for converting SASA to RSA.  Rose (Rose et al. 1985) Miller (Miller et al. 1987) Tien (Tien et al. 2013) |
| pdb_controls | use_rsa_sasa | “or”, “and” | Filtering logic for when both rsa_cutoff, and sasa_cutoff are provided |
| pdb_controls | only_exposed_in_patch | TRUE, FALSE | Provides the ability to have buried residues in windows, though buried residues will not be spatial window seeds |
| pdb_controls | interface_dist_cutoff | 5, numeric() | Angstrom distance between heavy atoms for defining interface contacts |
| pdb_controls | force_file_type | NULL, “pdb”, “cif” | Control which PDB file reading function to use from bio3d (Grant et al. 2006). Default NULL uses file extensions to determine how to read in PDB file. |
| aln_controls | use_sample_names | TRUE, FALSE | For cross-gene contexts will spatial haplotypes be built only across sequences with the same fasta header or across any sequences. When FALSE and MSA inputs have a different amount of sequences, the remaining sequences of the longer MSA are concatenated with gap residues |
| aln_controls | auto_chain_threshold | 0.2, numeric() | Kmer overlap threshold for determining MSA to PDB chain mappings |
| aln_controls | kmer_size | 4, numeric() | Kmer size for auto chain mapping |
| collapse_controls | merge_type | “exposure_distance”, “distance”, “union” | For merging per residue windows into per codon. Parameter covered in more detail in section **S.2** |
| collapse_controls | merge_exposure | 0.5, numeric() | Threshold for codon collapsed exposure, at 0.5 codons with 50% or more of their residue exposed (defined by user rsa_cutoff and sasa_cutoff) are classified as exposed and are seeds for spatial windows |
| stat_controls | calc_pi | FALSE, TRUE | Calculate nucleotide diversity with pegas package (Paradis 2010). Only for nucleotide MSA inputs |
| stat_controls | calc_tajima | FALSE, TRUE | Calculate Tajima’s D with pegas package (Paradis 2010). Only for nucleotide MSA inputs |
| stat_controls | calc_hap | FALSE, TRUE | Calculate Haplotype diversity with pegas package (Paradis 2010). Only for nucleotide MSA inputs |
| stat_controls | calc_site_entropy | TRUE, FALSE | Calculate amino acid Shannon entropy (Shannon 1948) in bits |
| stat_controls | calc_avg_patch_entropy | FALSE, TRUE | Calculate average single site Shannon entropy (Shannon 1948) per window |
| stat_controls | calc_block_entropy | FALSE, TRUE | Calculate amino acid haplotype Shannon entropy (Shannon 1948; Olsen et al. 2011) in bits |
| stat_controls | valid_aa_only | TRUE, FALSE | Controls excluding gap and non-noncanonical amino acid characters for the three entropy statistics |
| output_controls | output_dir | NULL, character() | Default NULL skips output writing entirely, otherwise a path is given |
| output_controls | write_msa_subsets | TRUE, FALSE | If output_dir is given spatial haploytpes are written as fasta files, one per spatial haplotype with file name corresponding to the msa_subset_id column in the final evo3d_df results data frame |
| output_controls | write_evo3d_df | TRUE, FALSE | If output_dir is given, write the results data frame as a tsv |
| output_controls | write_call_info | TRUE, FALSE | If output_dir is given, store defined parameter space, input file paths (if provided), and internal MSA to PDB mapping assignments for an analysis run as text file |
| output_controls | write_module_intermediates | TRUE, FALSE | If output_dir is given, store module intermediates like MSA to PDB alignments and PDB distance matrices, depending on the detail_level requested to run_evo3d() |
| output_controls | prefix | “”, character() | Prefix to append all file writes, if files with the same name exist, a timestamp code is prepended to all file writes to avoid overwriting data |

**S.2: Algorithmic definitions of codon-collapsing modes**

- **union** merges all residue-level windows corresponding to each codon, collecting every codon captured across those windows. Each codon is retained up to its maximum capture count observed in any single residue window, preserving within-window multiplicity. When max_patch is set, union overrides this parameter, and the resulting window sizes between codon-level windows can be different.
- **distance** merges residue-centred windows for each codon and serves as the core mode for generating fixed-size windows across residue environments. When max_patch = NA, distance behaves identically to union. When max_patch is defined, it constructs fixed-size windows that respect within-window exposure. To prevent oversampling of equivalent codons across multiple windows, quota masking is applied: once a codon is included, its nearest (first-ranked) instance is masked in all other residue-windows. This allows codon duplicates to be retained while preventing overcounting of shared neighbours. When patch_mode = "codon", codons are deduplicated, and distance continues filling until the fixed-size limit is reached or the dist_cutoff (if set) is exceeded.
- **exposure_distance** merges residue-centred windows for each codon using exposure-filtered geometry. When max_patch = NA, exposure_distance behaves like union but includes only codons meeting the global exposure threshold (e.g., exposure_threshold = 0.5, meaning a codon is considered exposed if at least half of its mapped residues are solvent-accessible). When max_patch is defined, it generates fixed-size patches from these exposure-qualified codons, continuing until the window reaches the specified count or the dist_cutoff (if set) is exhausted. The same minimum-distance and quota logic as distance is applied here.

**S.3: Implementation details and limitations of solvent-accessible surface area (SASA) computation** 
Validation against MKDSSP v4.4.10 (Hekkelman et al. 2025), as described in Methods, was performed after adjusting for two coordinate-handling differences between DSSP (Kabsch and Sander 1983; Hekkelman et al. 2025) and evo3D.

Within the run_evo3d() workflow, alternate atomic locations specifying multiple coordinates for a single atom are reduced to one representative position prior to SASA calculation. The MKDSSP v4.4.10 implementation includes all alternate locations when present, which can slightly alter solvent-accessible surface estimates. When SASA is computed directly using the standalone .calculate_accessibility() function, alternate locations may be provided.

Minor discrepancies may also arise from non-canonical amino acids that appear as HETATM records in the coordinate table but are listed in the SEQRES entry of the PDB or mmCIF file. The evo3D implementation relies on the PDB and mmCIF parsing functions of bio3D (Grant et al. 2006), which exclude HETATM residues from the sequence object. Converting such residues to ATOM records restores equivalence with DSSP outputs; however, within the wrapper function, these residues are omitted from both sequence and SASA calculations. 

**S.4: Notes on mixing MSA input types**
In the case of multiple MSA file inputs, three considerations arise. The first question, whether sequence names should be used to control spatial haplotype construction, has been covered in the main text and S.1.4 under use_sample_names. The second is MSA inputs having a different number of sequences. When use_sample_names is TRUE, this is not an issue as only sequences with shared names between all input MSAs are retained. However, when use_sample_names is FALSE, sequences are concatenated in the order they appear in the input files. The input file with more sequences is left with nothing to concatenate, so it is filled with gap characters. When valid_aa_only in the stats module is TRUE, no issues arise; but when set to FALSE, a considerable number of artificial gap characters can distort the statistics. Therefore, a poor parameterisation would be one with different sequence-count MSA inputs, use_sample_names set to FALSE, and valid_aa_only set to FALSE. The final consideration is the mixing of nucleotide and protein MSA inputs. This is valid, but the constructed spatial haplotypes will be forced to be amino-acid spatial haplotypes. Amino acid spatial haplotypes are limited, in the stats module, to the entropy set of predefined statistics, as no nucleotide statistics can be employed.

**S.5: General recommendations on patch_mode, analysis_mode, and patch type combinations**
The evo3D framework defines spatial windows that have not been used previously. Previous methods can be grouped into one parametrization: patch_mode = ‘residue’, analysis_mode = ‘residue’, distance patch types (set by dist_cutoff = … and max_patch = NA). Here, we have expanded both patch_mode and analysis_mode to include codon-defined, which follows three codon-merging schemes (**S.2**), and patch type to be fixed-count defined. The following table provides a brief recommendation on appropriate use.

Table S3: **General recommendations of appropriate use of patch_mode, analysis_mode, and patch type combinations**

| patch_mode | analysis_mode | patch_type | Appropriate use |
| --- | --- | --- | --- |
| residue | residue | distance | Haplotype statistics where codon duplications will not skew statistics (block entropy, nucleotide diversity, haplotype diversity), and desire to see diversity changes across different conformations of the protein system. A caveat is there needs to be a window size normalisation to accurately compare statistics across windows. |
| residue | residue | fixed count | Same as above, without needing window size normalisation of statistics |
| residue | codon  (merge_type = ‘union’) | distance | Haplotype statistics where codon duplications will not skew statistics. Need to normalize statistics by window size. union merging unifies all the spatial environments of this codon into one window. |
| residue | codon  (merge_type = ‘distance’) | distance | Algorithmically equivalent to above. distance and union merge types differ only for fixed count windows |
| residue | codon  (merge_type = ‘exposure_distance’) | distance | Haplotype statistics where codon duplications will not skew statistics. Need to normalize statistics by window size. This implementation reduces influence of codons that are usually buried but appear exposed in some PDB contexts. |
| residue | codon  (merge_type = ‘union’) | fixed count | Haplotype statistics where codon duplications will not skew statistics. Need to normalize statistics by window size. union merging here will result in variable count windows, it is the union of underlying fixed count windows. |
| residue | codon  (merge_type = ‘distance’) | fixed count | Haplotype statistics where codon duplications will not skew statistics. Provides fixed count windows per codon. |
| residue | codon  (merge_type = ‘exposure_distance’) | fixed count | Haplotype statistics where codon duplications will not skew statistics. Provides fixed count windows of residues whose codons pass the merge_exposure threshold. |
| codon | residue | distance | Haplotype or spatially averaged single site statistics, and desire to see diversity changes across different conformations of the protein system. A caveat is there needs to be a window size normalisation to accurately compare statistics across windows. |
| codon | residue | fixed count | Same as above, without needing window size normalisation of statistics |
| codon | codon  (merge_type = ‘union’) | distance | Haplotype or spatially averaged single site statistics. Need to normalize statistics by window size. union merging unifies all the spatial environments of this codon into one window. |
| codon | codon  (merge_type = ‘distance’) | distance | Algorithmically equivalent to above. distance and union merge types differ only for fixed count windows |
| codon | codon  (merge_type = ‘exposure_distance’) | distance | Haplotype or spatially averaged single site statistics. Need to normalize statistics by window size. This implementation reduces influence of codons that are usually buried but appear exposed in some PDB contexts. |
| codon | codon  (merge_type = ‘union’) | fixed count | Haplotype or spatially averaged single site statistics. Need to normalize statistics by window size. union merging here will result in variable count windows, it is the union of underlying fixed count windows. |
| codon | codon  (merge_type = ‘distance’) | fixed count | Haplotype or spatially averaged single site statistics. Provides fixed count windows per codon. |
| codon | codon  (merge_type = ‘exposure_distance’) | fixed count | Haplotype or spatially averaged single site statistics. Provides fixed count windows of residues whose codons pass the merge_exposure threshold. |

**S.6 Runtime and memory scaling of run_evo3d()**
Computational scaling of run_evo3d() was evaluated using the Chikungunya virus E1/E2 protein complex, PDB ID: 8fcg (Chmielewski et al. 2024). Following the use-case examples presented in the manuscript, pdb_controls were set to sasa_cutoff ≥ 10 and max_patch = 15, and distance_method was varied between "centroid" and "all". No downstream population genetic statistics were applied; therefore, this benchmark reflects only the time and memory required for MSA-to-PDB mapping, spatial window construction, and extraction of spatial haplotypes returned to the user.

The test was performed on four subsets of the structural model, increasing from one chain of each gene, chain = (‘A’, ‘E’), to four chains of each gene, chain = (‘ABCD’, ‘EFGH’). Runtime and peak memory usage were recorded. Each configuration was executed twice, and the reported values represent the average runtime and memory usage. Runtime was measured using system.time() and peak memory usage using the peakRAM R package (Quinn 2017).

**Table S4. Runtime and peak memory usage of run_evo3d() across increasing structural sizes of the Chikungunya E1/E2 complex (PDB ID 8fcg).** Runtime in seconds (s) and peak memory usage (MiB) are reported for the "all" pairwise atom distance and the "centroid" distance methods. Values represent the mean of two runs for each structural subset.

| Residues | Atoms | Spatial windows | Runtime (s) all | Runtime (s) centroid | Peak memory (MiB) all | Peak memory (MiB) centroid |
| --- | --- | --- | --- | --- | --- | --- |
| 858 | 6622 | 653 | 7.7 | 5.6 | 805.0 | 250.3 |
| 1716 | 13244 | 1298 | 15.9 | 7.5 | 3278.8 | 389.4 |
| 2574 | 19866 | 1925 | 30.6 | 10.3 | 6789.5 | 442.2 |
| 3432 | 26488 | 2573 | 91.2 | 13.8 | 11076.5 | 558.7 |

**S.7 Post-analysis functions**

**S.7.1: generating analysis matched null models**
After analysis, the full results list can be passed to generate_null_model(), which constructs a specified number of synthetic spatial haplotypes from the codon pool included in the analysis spatial windows. Two modes of codon-matched analysis are available, controlled by match_codon_frequency, sampling codons at the frequency they are included in analysis windows, or sampling uniformly for all codons included in the analysis windows. An example generating 10,000 synthetic subsets of the MSAs, all containing 15 codons, follows:

```
results = run_evo3d(...)
null_haplotypes = generate_null_model(
	results, n = 10000, len = 15, seed = 123
)
```

**S.7.2: filtering overlapping windows into reduced sets**
For visualisation or statistical testing, it may be useful to reduce the set of spatial windows. filter_overlaps() performs pruning in the row order of an input evo3d_df. The parameter, overlap, controls the shared codon allowance for a window and the windows already included in the reduced set. Filtering for less than 33% overlap follows, continuing with the example from **S.7.1**:

evo3d_df = results$evo3d_df
sorted_df = evo3d_df[order(evo3d_df$block_entropy), ]
reduced_df = filter_overlaps(sorted_df, overlap = 0.33)

**S.7.3: Writing statistics to PDB files for visualisation**
Examining results in PyMol (Schrödinger and DeLano 2020) or other molecular-visualisation software is aided by write_stat_to_pdb(). The full results list is passed to the function, with additional arguments controlling the statistic column to add to the b-factor and occupancy columns, replacement value for NA values, analysis PDB to map values, and whether the output PDB file is to contain all original chains or only analysis chains. When multiple column names are passed to stat_name the first replaces b-factor and the second occupancy. An example continuing with **S.7.2**:

write_stat_to_pdb(
results, pdb_id = ‘pdb1’,
stat_name = ‘block_entropy’,
adjust_NA_stats = -99,
mapped_chains_only = TRUE,
outfile = ‘block_entropy.pdb’

)

With PyMol command structure as:

load block_entropy.pdb
select stat, b > -99
spectrum b, selection=stat

**Supplementary Figures**

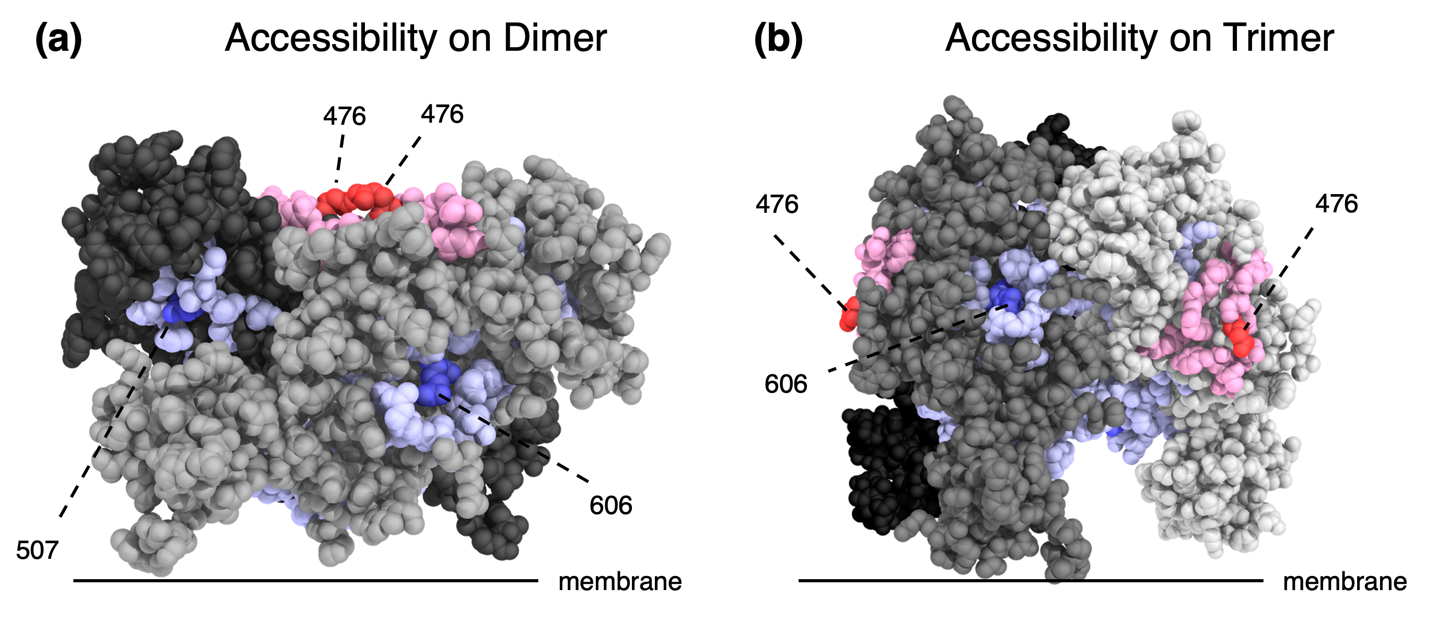

**Figure S1. Accessibility of significantly conserved and diverse surface patches in the Hepatitis C virus E1E2 complex.** **(a)** Significantly conserved (blue) and diverse (red) neighbourhoods identified in Fig. 2c (PDB ID: 8fsj) are shown superimposed on the experimentally solved E1E2 dimer (PDB ID: 8rjj; RMSD = 1.295 across 2043 atoms). The neighbourhood of residue 606 is surface-accessible, while the neighbourhood of residue 507 is likely less accessible as an epitope due to steric hindrance. The neighbourhood of residue 476 is diverse and fully accessible, consistent with known frequent immune recognition. **(b)** The same significant patches mapped to the predicted trimer by superimposing 8fsj onto an AlphaFold3-predicted trimer (RMSD = 2.091 across 2176 atoms). Neighbourhoods for residues 606 and 476 are predicted to be accessible, while the conserved neighbourhood around residue 507 participates in the predicted trimer interface.

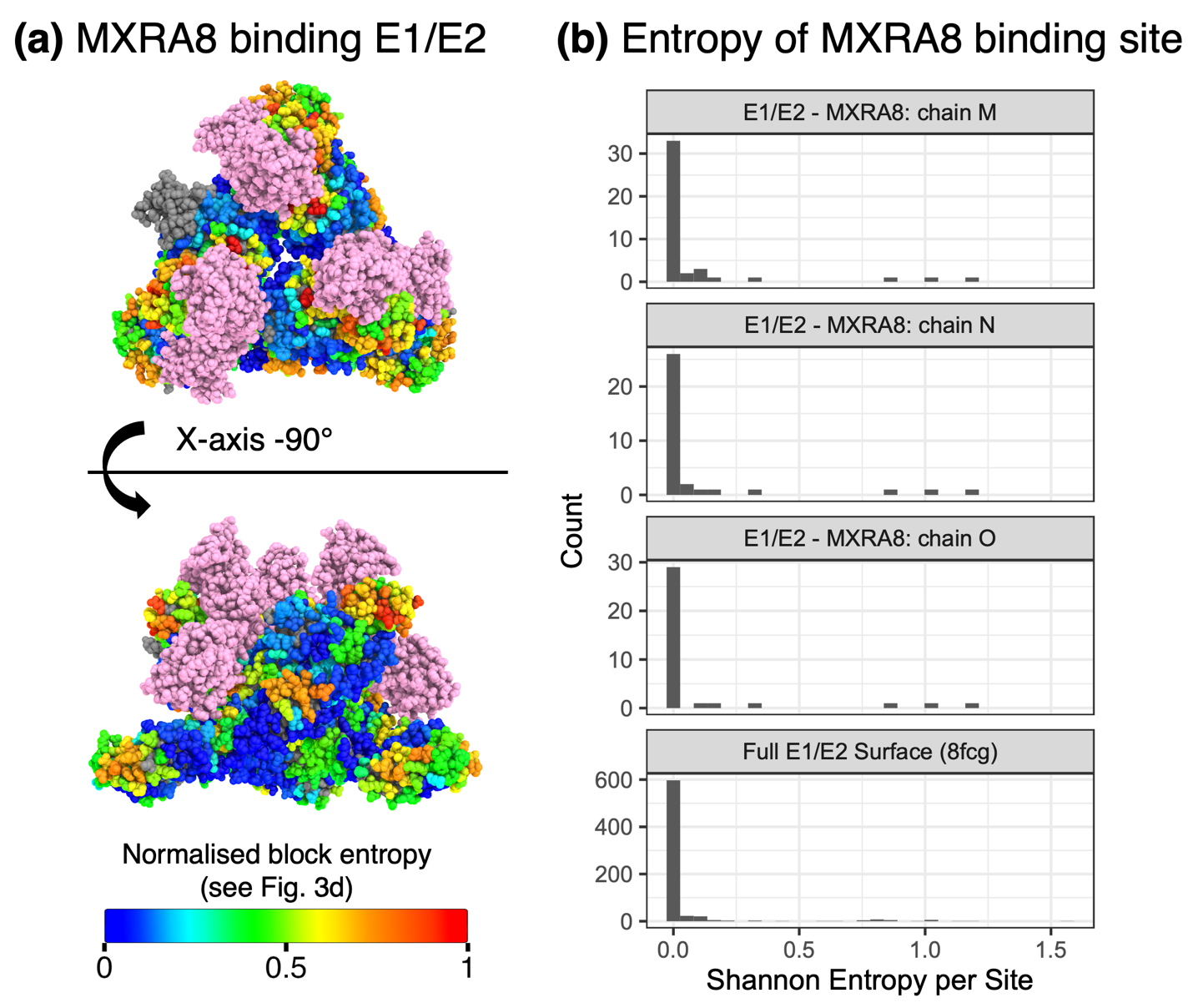

**Figure S2. Interface-aware capture in evo3D enables binding interfaces to be analysed as spatial neighbourhoods. (a)** MXRA8 (pink) bound to the Chikungunya virus E1/E2 complex (PDB 6jo8), with the viral surface coloured by normalised block-entropy values derived from spatial windows constructed on the octameric assembly (PDB 8fcg) (see Fig. 3d). This visualisation shows how variable and conserved spatial patches identified in Fig. 3d relate to the MXRA8-binding footprint. The grey region in the upper left corresponds to the E3 protein, for which no MSA information was generated in this study. **(b)** Per-site amino acid entropy across the three copies of E1/E2 - MXRA8 binding interface compared to per-site entropy of the full E1/E2 surface (computed using 8fcg). Interface contacts were defined from 6jo8. Because both 6jo8 and 8fcg map to the same MSAs, codon positions are directly comparable, allowing an interface defined in one structure to be evaluated on the other.
